# Quality control-based signal drift correction and interpretations of metabolomics/proteomics data using random forest regression

**DOI:** 10.1101/253583

**Authors:** Hemi Luan, Fenfen Ji, Yu Chen, Zongwei Cai

**Author notes:** These authors contributed equally to this work (Hemi Luan, Fenfen Ji).

## Abstract

Large-scale mass spectrometry-based metabolomics and proteomics study requires the long-term analysis of multiple batches of biological samples, which often accompanied with significant signal drift and various inter‐ and intra‐ batch variations. The unwanted variations can lead to poor inter‐ and intra-day reproducibility, which is a hindrance to discover real significance. We developed a novel quality control-based random forest signal correction algorithm, being ensemble learning approach to remove inter‐ and intra‐ batches of unwanted variations at feature-level. Our evaluation based on real samples showed the developed algorithm improved the data precision and statistical accuracy for metabolomics and proteomics, which was superior to other common correction methods. We have been able to improve its performance for interpretations of large-scale metabolomics and proteomics data, and to allow the improvement of the data precision for uncovering the real biologically differences.

## INTRODUCTION

Mass spectrometry (MS)-based omics techniques, including metabolomics and proteomics, are rapidly growing fields in system biology combining both analytical and statistical methodologies for a high throughput analysis of multiplex molecule profiles. Large-scale experiments coupled with varieties of statistical tools have been used to identify specific biological changes, leading to the understanding of biomarkers and multi-biochemical pathways ^1,2^. However, the omics data obtained from MS-based experiments is subject to various forms of unwanted variations including both within-batch and between-batch variations introduced by signal drift/attenuation and multiplicative noise ^3^ (e.g., temperature changes within the instrument, accumulated contamination, or loss of instrument performance during a long run of samples). These inherent biases and variations in metabolomics and proteomics data were challenges for quantitative comparative analysis ^3,4^.

Quality control (QC) has been considered as an essential step in the large-scale and long-term biological study, allowing to evaluate the response signal drift, mass accuracy and retention time in intra‐ and inter-batch experiments ^5^. Based on a controlled experiment involved QC samples throughout the data collection process, the most advantageous uses of QC samples can be obtained, allowing the correction of signal drift and other systematic noise through mathematical algorithms in liquid or gas chromatography (LC or GC) hyphenated to MS-based metabolomics study ^5,6^. However, there is few such controlled experiment and algorithms for improving the data quality in label-free LC-MS based proteomics ^4,7^. Given that technological advances in QC-based experimental designs and the growth of large-scale metabolomics and proteomics projects, as well as the requirement of the accuracy and efficiency for handling unwanted variations, the new computational algorithms support is highly demanded. Therefore, we developed a novel QC-based random forest signal correction (QC-RFSC) algorithm to remove unwanted variations at feature-level in large-scale metabolomics and proteomics data.

## MATERIALS and METHODS

### Feature-based signal drift correction

In the following, for any feature j in any sample i, the intensity value in the preprocessed data is represented by x_ij_ and the corrected value by x’_ij_. Due to the different degree of drift in each feature j in a sample i, the correction factor F_ij_ was assigned according to eq 1.

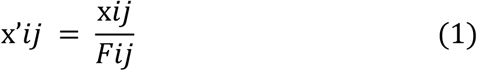

The correction factor of each feature was obtained by learning an ensemble of regression trees each grown on a random sub - QC sample to infer a noisy variant during the experimental analysis, and calculated using the regression fit derived from the intensities of the QC injections. For a feature j in the QC-RFSC method, each sample i will have a different F_ij_ according to its position k_ij_ in the injection order as defined by eq 2:

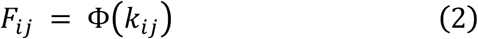

Φ represents random forests model with 500 regression trees. After QC-RFSC procedure and batches integration, the low quality of peaks could be filtered using quality assessment criteria ^2^. Two additional common signal drift correction methods, such as QC-RLSC ^2^ and QC-SVR ^8^ were also tested in our study. The correction factors in QC-RLSC and QC-SVR methods were calculated using the nonlinear local polynomial regression (LOESS) and support vector regression (SVR) fit derived from the intensities of the QC injections, respectively.

### Sample preparation and LC-MS analysis

Urine preparation for metabolomics analysis was performed as previously reported^9^. A Dionex U3000 LC system coupled online to Q Exactive Focus instrument (Thermo Fisher Scientific, MA, USA) was employed to detected the urinary metabolic profile. On the other hand, serum preparation including protein denaturation, digestion, and alkylation in proteomics analysis were performed as previously reported ^10^. Total of 174 transitions response to 69 proteins were monitored by using a Dionex U3000 LC system coupled online to TSQ Quantiva instrument (Thermo Fisher Scientific, MA, USA) with selected reaction monitoring (SRM) mode. The details of the analytical experiments were described in supplemental materials.

### Study designs

In this work, two non-targeted metabolomics and one quantitative SRM-based proteomics studies were involved to evaluate the performance of QC-RFSC method and statTarget 2.0 software.

Firstly, we performed a MS-based metabolomics experiment for the investigation of the urinary metabolic profiles (Dataset 1). One pooled urine sample and another individual urine sample were used as the QC sample and real sample, respectively. The aliquoted QC samples were analyzed after every five aliquoted real samples in the entire batch. In total, we produced 4549 metabolic features, which were consistently detected in 46 QC samples and 175 aliquoted urine samples. Secondly, we collected a standard MS-based metabolomics data (Dataset 2) from GigaScience databases (ID: 100108) ^11,12^. A total of individual 180 pregnant women were recruited and divided into six groups (A-F) according to their gestational weeks. The aliquoted 38 QC samples were analyzed after every six aliquoted real plasma samples in the two batches. The detailed description of Dataset 2 was provided in supplemental materials. Thirdly, we designed a SRM-based proteomics study for quantitative analysis of protein markers of Parkinson’s disease (PD). The study was approved by the ethics committee of Hong Kong Baptist University’s Institutional Review Board, and written informed consents were obtained in this study. A total of 34 PD patients together with 34 normal control subjects were recruited. Venous blood was collected with blood collection tubes in the morning before breakfast from all the participants, and then serum samples were separated at room temperature from vein blood and stored at −80 °C until use. The pooled QC samples were analyzed by LC-MS after every eight subject’s samples in the entire batch. The running order of subjects was also randomized. This study was used to evaluate the performance of the QC-RFSC method for removal of inter-batch variations in the proteomics data (Dataset 3).

### Data analysis

MS raw data files were converted to mzXML format using ProteoWizard. The XCMS^13^ and Skyline^14^ was used for the extraction of peak abundances of metabolites and tryptic peptides, respectively. The statTarget was employed for signal drift correction at feature-level and statistical analysis.

## RESULTS and DISCUSSION

### Signal drift correction

Supervised machine learning allows for systematic pattern identification from a high-dimensional features in the training dataset and minimizes manual tuning for optimal model generalization. It is well-suited for an unbiased analysis of MS data, and have been used for identification of metabolite fingerprinting, biomarkers and protein sequences. Twelve well-known supervised machine learning algorithms were evaluated by using the caret package with 10-fold cross validation ^15^, such as random forest (RF), bayesian, eXtreme gradient boosting, support vector machines, LOESS, linear regression, etc. Our results showed the RF, QRF and Loess outperformed with high predictive accuracy or R-square at median and first quartile value for the dataset 1 (**Fig. 1**).

**Figure 1.**
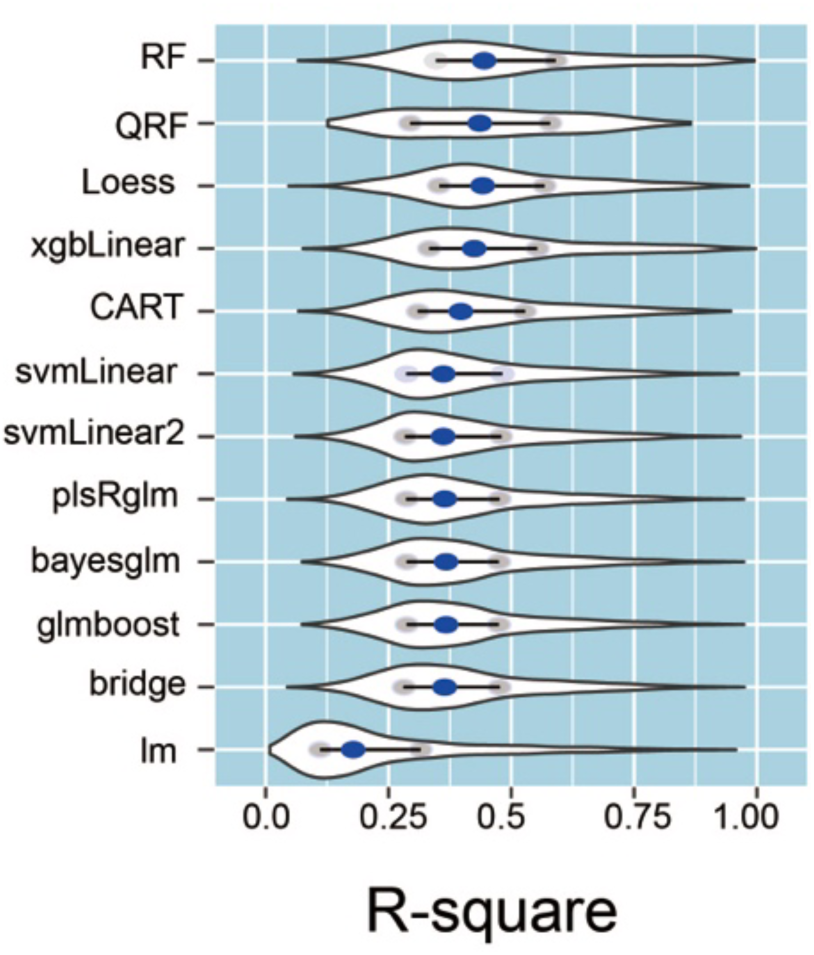
Comparison of the performance of machine learning models. Violin plot of the predictive accuracy of twelve machine learning models on dataset 1. Two grey dots denotes the first and third quantile, respectively; Blue dot, median value;

Given that RF performed well in the real dataset and being one of the most accurate learning algorithms available for MS-based omics data, RF was therefore selected for QC-based signal correction ^16,17^. QC-RFSC algorithm integrates the RF based ensemble learning approach to learn the unwanted variations from QC samples, and predict the correction factor in the neighboring real samples responses. To evaluate the performance of QC-RFSC algorithms, we compared available algorithms that were used for QC-based signal correction, including RF, SVR ^8^ and LOESS ^2^. We further calculated the cumulative frequency of RSD% of all features (**Fig. 2**). In the raw data, the proportion of peaks within 15 % RSDs was only 1.97 % of the total peaks. After adjusted by QC-RFSC method, there were a 12.7-fold increase (25.1%) in the number of peaks within 15 % RSDs. Meanwhile, the proportion of peaks with RSDs less than 30 % increased significantly from 43.9% to 86.2 % after QC-RFSC method. The QC-RLSC and QC-SVR algorithms also increased the proportion of peaks with 30% RSDs to 65.5% and 61.9 %, respectively (**Fig. 2**). Our results demonstrated that the QC-RFSC robustly increased precision of metabolomics data and statistical accuracy through removal of unwanted variations, which had better performance than the other two methods.

**Figure 2.**
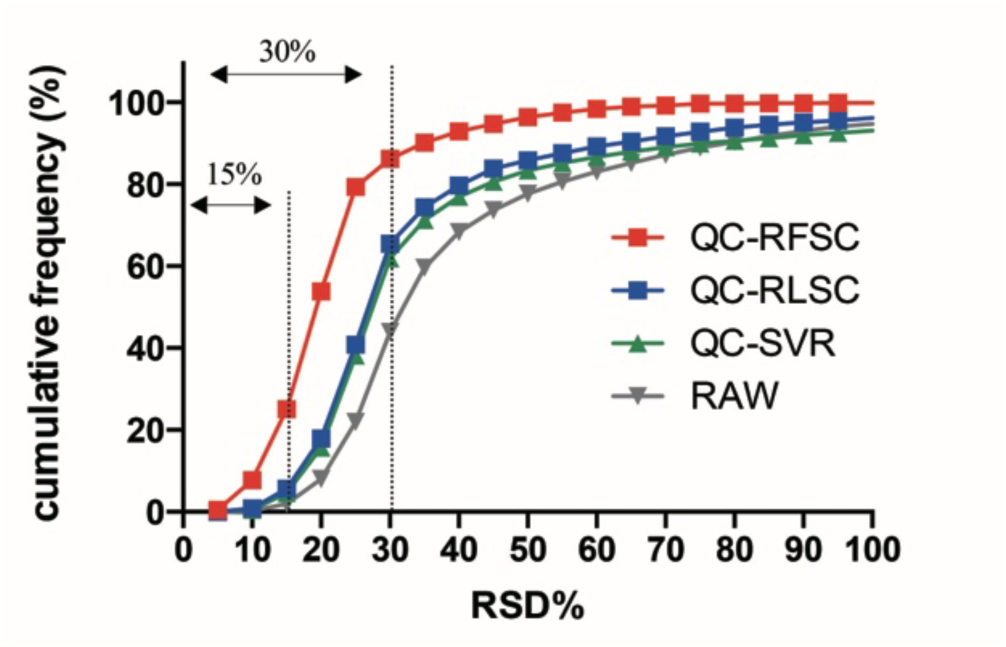
Comparison of the cumulative frequency of RSD% of all features with three correction methods

### Precision improvement for metabolomics and proteomics data

Precision is one of the most important criteria in the assessment of an analytical method, and achieved by monitoring quality control samples during analysis^5^. The percentage RSD% of each peak in QC samples was usually used for precision evaluation. To evaluate the precision improvement that can be achieved by our software, we evaluated the cumulative RSD distribution of all features in metabolomics data (dataset 2) and proteomics data (dataset 3) with pre‐ and post‐ QC-RFSC, QC-RLSC and QC-SVR (**Fig. 3-4**). As a comparison, the RSDs of peaks in metabolomics data were significantly decreased across the entire range using the QC-RFSC, QC-RLSC and QC-SVR methods (**Fig. 3A**). The proportion of peaks with RSDs less than 30 % increased significantly to 90.9% after QC-RFSC compared with the raw data (52.1%). The QC-RLSC and QC-SVR also increased the proportion of peaks within 30% RSDs to 80.1% and 72.3%, respectively. The PCA score plots showed the QC samples were tightly clustered due to QC-RFSC correction (**Fig. 3B, C**).

**Figure 3.**
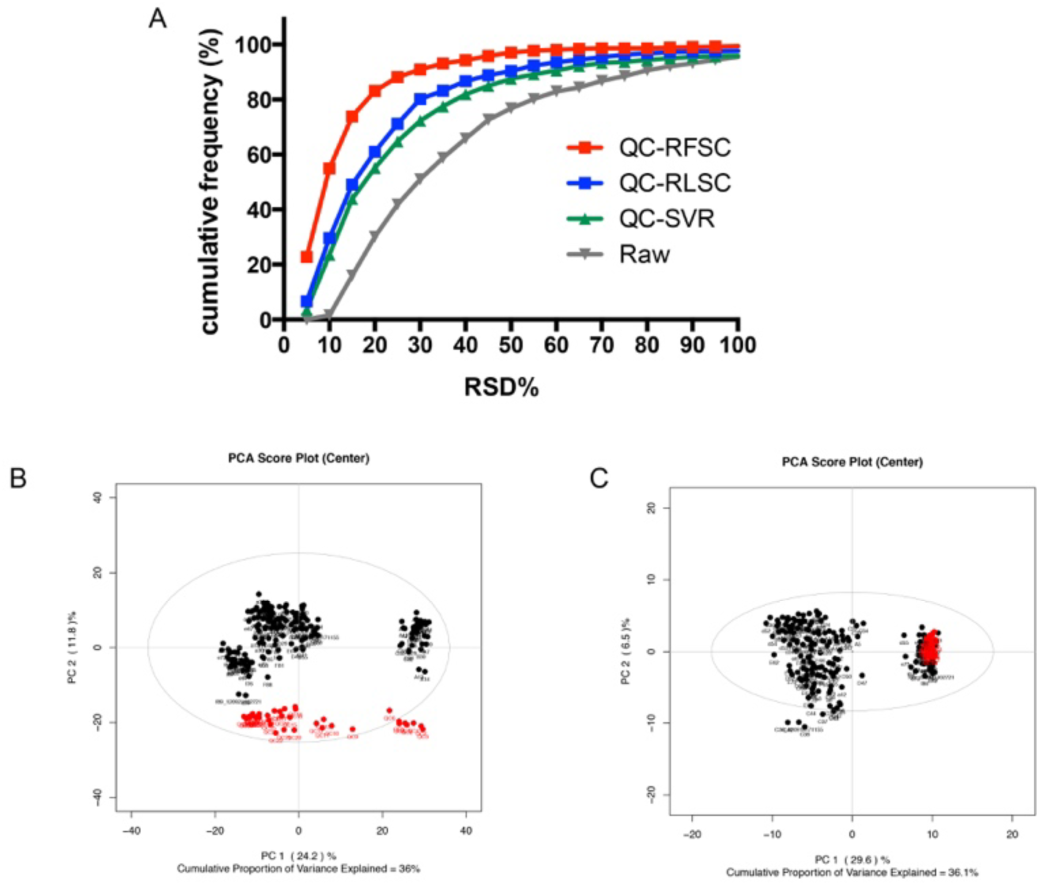
The performance of QC-RFSC method for metabolomics data. A, comparison of the cumulative frequency of RSD% of features in metabolomics data (dataset 2). B and C, PCA score plots of the metabolomics data with pre‐ (B) and post‐ (C) correction. Red dots denote the QC samples; Black dot, real samples.

**Figure 4.**
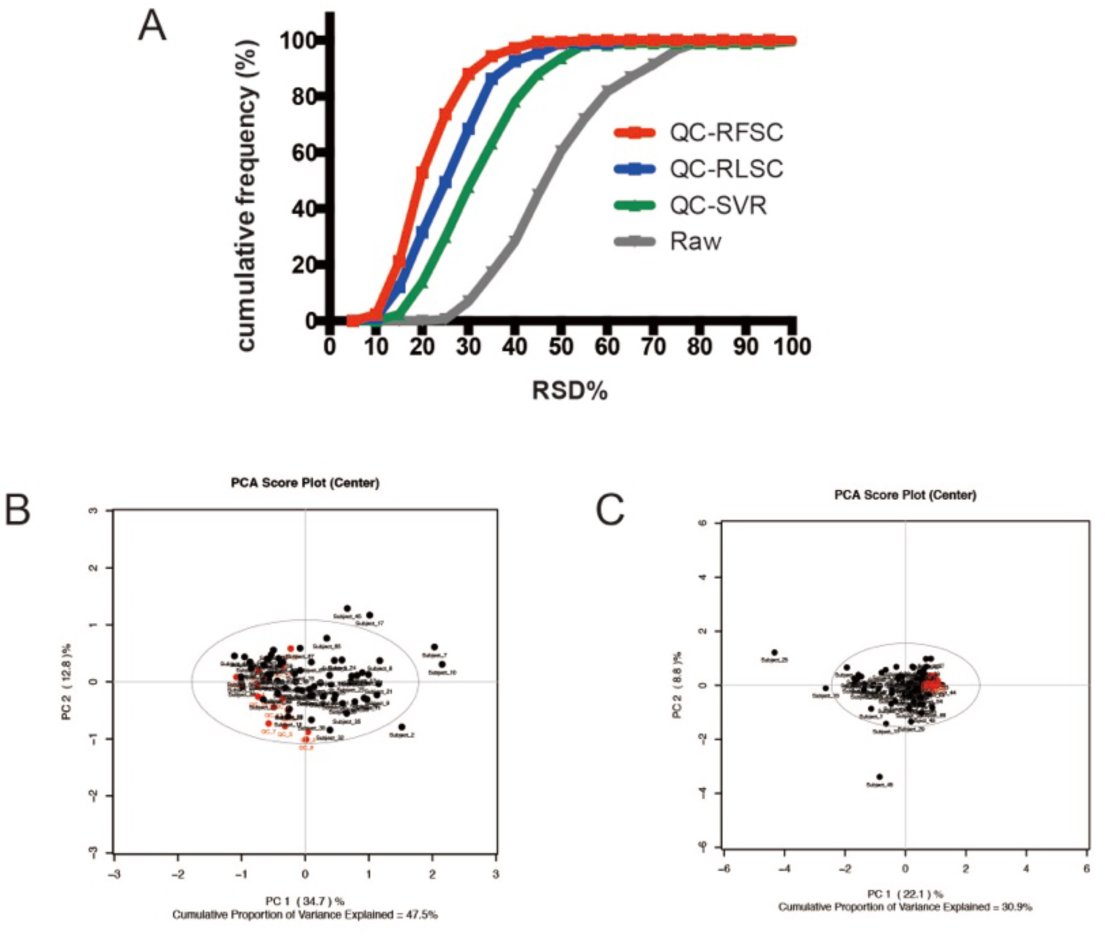
The performance of QC-RFSC method for proteomics data. A, comparison of the cumulative frequency of RSD% of features in proteomics data (dataset 2). B and C, PCA score plots of the proteomics data with pre‐ (B) and post‐ (C) correction. Red dots denote the QC samples; Black dot, real samples.

Beside metabolomics data analysis, the QC-RFSC could also significantly improve the data quality for the proteomics. The proportion of peaks within 30 % RSDs increased significantly to 87.9% after QC-RFSC compared with the QC-RLSC (68.4%) and QC-SVR (47.7%) (**Fig. 4A**). The clustered QC samples in PCA score plots indicated the data precision was significant improved. (**Fig. 4B, C**). The results demonstrated that the QC-RFSC significantly increased precision of metabolomics and proteomics data and had better performance than the two other methods.

It is noted that label-free proteomics experiments were much more complex than metabolomics experiments ^18^. The complicated sample preparation procedures (i.e. pre-fraction, digestion, desalting) may produce additional variations caused by systematic or random errors ^4,18^. Our results showed that the QC-RFSC is an efficient method to remove inter‐ and intra‐ batch of unwanted variations at feature-level, and improve the data precision and statistical accuracy for metabolomics and targeted proteomics.

## CONCLUSION

The developed QC-RFSC algorithm is a highly efficient approach to remove unwanted variations, to improve the data quality of metabolomics and proteomics data, and to further enlarge the number of differentially expressed features. Due to the substantial similarities among different types of expression data from system biology analysis (e.g., high dimension, analytical bias, significance analysis and so on), it was also feasible to extend the application of QC-RFSC algorithm from quantitative metabolomics data to other system biology data such as protein or peptide expression data.

## ACKNOWLEDGMENTS

The authors would like to thank the financial supports from Hong Kong Baptist University (IRMC/13-14/03-CHE) and the National Sciences Foundation of China (NSFC21675176 and NSFC91543202).

